# Exploring the potential genetic heterogeneity in the incidence of hoof and leg disorders in Austrian Fleckvieh and Braunvieh cattle

**DOI:** 10.1101/2020.01.11.902767

**Authors:** Barbara Kosinska-Selbi, Tomasz Suchocki, Christa Egger-Danner, Hermann Schwarzenbacher, Magdalena Fraszczak, Joanna Szyda

## Abstract

**Background:** Genetic heterogeneity denotes the situation when different genetic architectures underlying diverse populations result in the same phenotype. In this study, we explore the nature of differences in the incidence of the number of hoof and leg disorders between Braunvieh and Fleckvieh cattle in the context of genetic heterogeneity between the breeds.

**Results:** Despite potentially higher power of testing due to twice as large sample size, none of the SNPs was significantly associated with the number of hoof and leg disorders in Fleckvieh, while 16 SNPs were significant in Braunvieh. The most promising candidate genes in Braunvieh are: CBLB on BTA01, which causes arthritis in rats; CAV2 on BTA04, which in effects mouse skeletal muscles; PTHLH on BTA05, which causes disease phenotypes related to the skeleton in humans, mice and zebrafish; SORCS2 on BTA06, which causes decreased susceptibility to injury in the mouse. Some of the significant SNPs (BTA01, BTA04, BTA05, BTA13, BTA16) reveal allelic heterogeneity – i.e. differences due to different allele frequencies between Fleckvieh and Braunvieh. Some of the significant regions (BTA01, BTA05, BTA13, BTA16) correlate to inter-breed differences in LD structure and may thus represent false-positive heterogeneity. However, positions on BTA06 (SORCS2), BTA14 and BTA24 mark Braunvieh-specific regions.

**Conclusions:** We hypothesise that the observed genetic heterogeneity of hoof and leg disorders is a by-product of multigenerational differential selection of the breeds – towards dairy production in the case of Braunvieh and towards beef production in the case of Fleckvieh. Based on the current data set it is no possibly to unequivocally confirm/exclude the hypothesis of genetic heterogeneity in the susceptibility to leg disorders between Fleckvieh and Braunvieh because only explore it through associations and not the causal mutations. Rationales against genetic heterogeneity comprise a limited power of detection of true associations as well as differences in the length of LD blocks and in linkage phase between breeds. On the other hand, multigenerational differential selection of the breeds and no systematic differences in LD structure between the breeds favour the heterogeneity hypothesis at some of the significant sites.

## Background

Genetic heterogeneity denotes the situation when different genetic architectures underlying diverse populations result in the same phenotype. In human genetics, for decades, the concept of genetic heterogeneity has been considered in genome-wide association studies (GWAS) [1]. One of the most well-known diseases characterised by high degree of genetic heterogeneity is the human autism spectrum disorder [2]. Relatively recently the concept of genetic heterogeneity has also been introduced into to the analysis of data from artificially selected plant and livestock species by Bérénos et al. [3], de los Campos et al. [4], and Lehermeier et al. [5]. In such species, an important source of genetic heterogeneity may be due to a complex population structure, which is typically composed of divergently selected breeds exhibiting high variation in allele frequencies and linkage disequilibrium patterns [6].

Cattle hoof and leg disorders are relatively novel traits represented by a group of different phenotypes varying from binary, directly assessed disease diagnoses like e.g. a sole ulcer to composite traits scored on a categorical basis, e.g. a locomotion score. Due to their impact on welfare, productivity and fertility [7], the traits rapidly gain importance in dairy cattle breeding schemes. Technically, a common feature is a relatively poor definition of traits from this group and lack of routine recording, resulting in a large number of relatively small data sets scattered across various populations. Those features not only cause low power of detection of significant gene (or SNP) – phenotype associations resulting in a low reproducibility of results [7], but also imply a potential heterogeneity in the genetic determination of phenotypes due to differences in selection schemes and thus underlying differences in linkage disequilibrium and allele frequencies between populations [8].

In this study, we explore the nature of differences in the incidence of the number of hoof and leg disorders between Austrian bred Braunvieh and Fleckvieh cattle in the context of genetic heterogeneity between the breeds.

## Results

### Heterogeneity in association signals

Adapting the false discovery rate (FDR) threshold of 10%, despite potentially higher power of testing due to twice as large sample size, none of the SNPs was significantly associated with the number of hoof and leg disorders in Fleckvieh, while 16 SNPs were significant in Braunvieh (Figure 1). One of the three significant SNPs from BTA01 is located 285,955 bp upstream of CBLB gene, known to cause arthritis in rats. The same SNP is located within a region of a QTL for hindquarter proportions. The two most significant SNPs were located on BTA04. Both were intergenic, but their closest downstream gene encodes caveolin 2 protein, which in the mouse is known to effect skeletal muscles. A SNP on BTA05 falls within four QTL regions responsible for rump conformation traits. One of the most interesting significant annotation points at another SNP on BTA05, which is located 76,362 bp downstream of PTHLH. In humans and mice, this gene causes multiple disease phenotypes related to the skeleton. In zebrafish mutations of this gene result in decreased bone mineralisation, in humans – to brachydactyly and to numerous bone and calcium related disease phenotypes in the mouse, including: decreased length of long bones, premature bone ossification, and increased osteocyte apoptosis. Another interesting significant annotation points at the intron of SORCS2 gene on BTA06, which was assigned to decreased susceptibility to injury phenotype in the mouse. The effect on muscles, albeit in Zebrafish, was assigned to the protein encoded by PIP4K2A, which is located close to the significant SNP on BTA13. The same SNP is also within a QTL region for rump angle. On BTA16, a significant association points out at ENSBTAG00000009943 involved in inflammatory response. Significant SNPs are summarised in Table 1 except a SNPs on BTA06, which could not be placed on the current reference assembly (ARS-UCD1.2).

**Table 1.** SNPs significant in BSW, based on the FDR≤0.10 threshold. SNP genomic information corresponds to the ARS-UCD1.2 reference genome.

| Position name | Closest Gene(s) | QTL | Effect | Increasing allele | FDR – effect | FDR – distance | S | P – frequency ratio |
| --- | --- | --- | --- | --- | --- | --- | --- | --- |
| 1:3,303,269<br>rs110488513 | Intergenic between<br>MIS18A and HUNK |  | 0.018 | A | 0.000008 | 1.0 | 0.163591 | 0.391039 |
| 1:43,542,488<br>rs41661497 | Intergenic between<br>DCBLD2 and COL8A1 |  | 0.012 | G | 0.000106 | 1.0 | 0.152780 | 0.229225 |
| 1:50,767,507<br>rs110811919 | Intergenic upstream of<br>CBLB | Hindquarter proportions<br>(7124) | 0.007 | G | 0.015146 | 1.0 | 0.152780 | 0.007376 |
| 4:52,028,036<br>rs110514562 | Intergenic between<br>CAV2 and TES |  | 0.029 | A | <10<sup>-6</sup> | 1.0 | 0.002094 | <10<sup>-7</sup> |
| 4:52,079,221<br>rs137336750 | Intergenic between<br>CAV2 and TES |  | 0.029 | G | <10<sup>-6</sup> | 1.0 | 0.002094 | <10<sup>-7</sup> |
| 5:61,220,624<br>rs41590733 | Intergenic upstream of<br>NEDD1 | Rump conformation<br>(3422, 3424, 1563,<br>20622) | 0.010 | A | 0.052989 | 1.0 | 0.005398 | 0.030671 |
| 5:81,769,685<br>rs109268584 | Intergenic between<br>CCDC91 and PTHLH |  | 0.008 | A | 0.043017 | 1.0 | 0.005398 | 0.008493 |
| 6:114,116,280<br>rs110962969 | Intron of<br>SORCS2 |  | 0.010 | C | 0.002456 | 1.0 | 0.006265 | 0.381230 |
| 13:13,590,662<br>rs110792762 | Intergenic upstream of<br>CELF2 |  | 0.008 | G | 0.012588 | 1.0 | 0.451395 | 0.003215 |
| 13:23,590,146<br>rs110989397 | Intergenic between<br>SPAG6 and PIP4K2A | Rump angle (3429) | 0.019 | A | <10<sup>-6</sup> | 1.0 | 0.006076 | 0.019040 |
| 14:55,768,446<br>rs110534995 | Intergenic between<br>TMEM74 and EMC2 |  | 0.010 | A | 0.003962 | 1.0 | 0.023641 | 0.153087 |
| 16:12,125,227<br>rs29024589 | Intergenic between<br>CDC73 and GLRX2 |  | 0.010 | T | 0.000676 | 1.0 | 0.021476 | 0.394140 |
| 16:12,280,122<br>rs110843300 | Intergenic upstream of<br>UCHL5 |  | 0.008 | G | 0.006398 | 1.0 | 0.021475 | 0.398872 |
| 16:36,037,389<br>rs41579631 | Intergenig between<br>RGS7 and<br>ENSBTAG00000009943 |  | 0.007 | G | 0.000592 | 1.0 | 0.026922 | <10<sup>-115</sup> |
| 24:24,273,191<br>rs136424124 | Intergenic between<br>CCDC178 and KLHL14 |  | 0.002 | A | 0.082299 | 1.0 | 0.006277 | 0.111094 |

**Figure 1.**
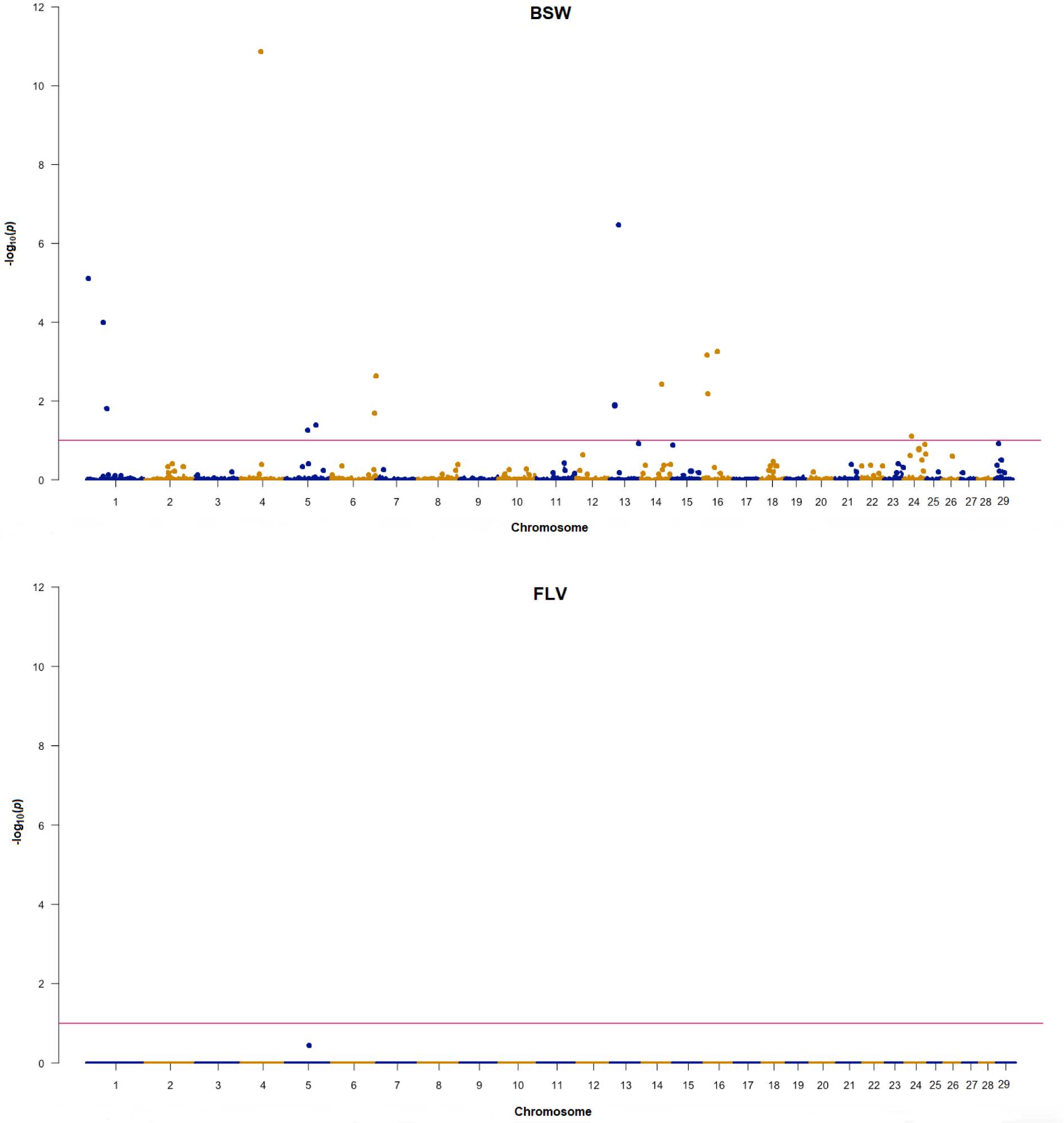
Manhattan plots for GWAS for hoof and leg disorders in Braunvieh and Fleckvieh. The horizontal line corresponds to FDR of 0.01.

There was no correlation between P values observed for SNPs in FLV and BSW, which was estimated to 0.00302 for all SNPs and −0.01494 (−0.08065) for SNPs with 100 smallest P values in BSW (FLV). In addition, breed-specific SNP effect estimates also revealed a very low correlation of 0.02363.

### Heterogeneity in genetic architecture

For the chromosomes containing SNPs significant in Braunvieh, Machalanobis distances, expressing differences between breeds in SNP genotype variability, and the S statistics, expressing differences in the LD decay pattern, were visualised on Figure 2. For the Machalanobis distance, all FDR values at the significant SNP locations are equal to one. Even while considering whole chromosomes harbouring significant SNPs none of the distances was significant, indicating that it was not possible to differentiate between breeds based on SNP genotypes corresponding to the 50-SNP windows. A somewhat different picture emerged when inter-breed differences in LD were considered. In some, but not all, of the regions, significant SNPs correspond to windows for which a difference ins LD structure was indicated by high values of the S statistics - rs110488513 and rs41661497 on BTA01 as well as rs110792762 on BTA13. Some other significant SNPs are located in windows adjacent to such windows - rs110811919 on BTA1, as well as rs29024589, rs110843300, and rs41579631 on BTA16. For eight significant SNPs (rs29024589 on BTA01, both SNPs on BTA04, both SNPs on BTA05, both SNPs on BTA13, rs41579631 on BTA16) significant allelic heterogeneity was detected (Table 1).

**Figure 2.**
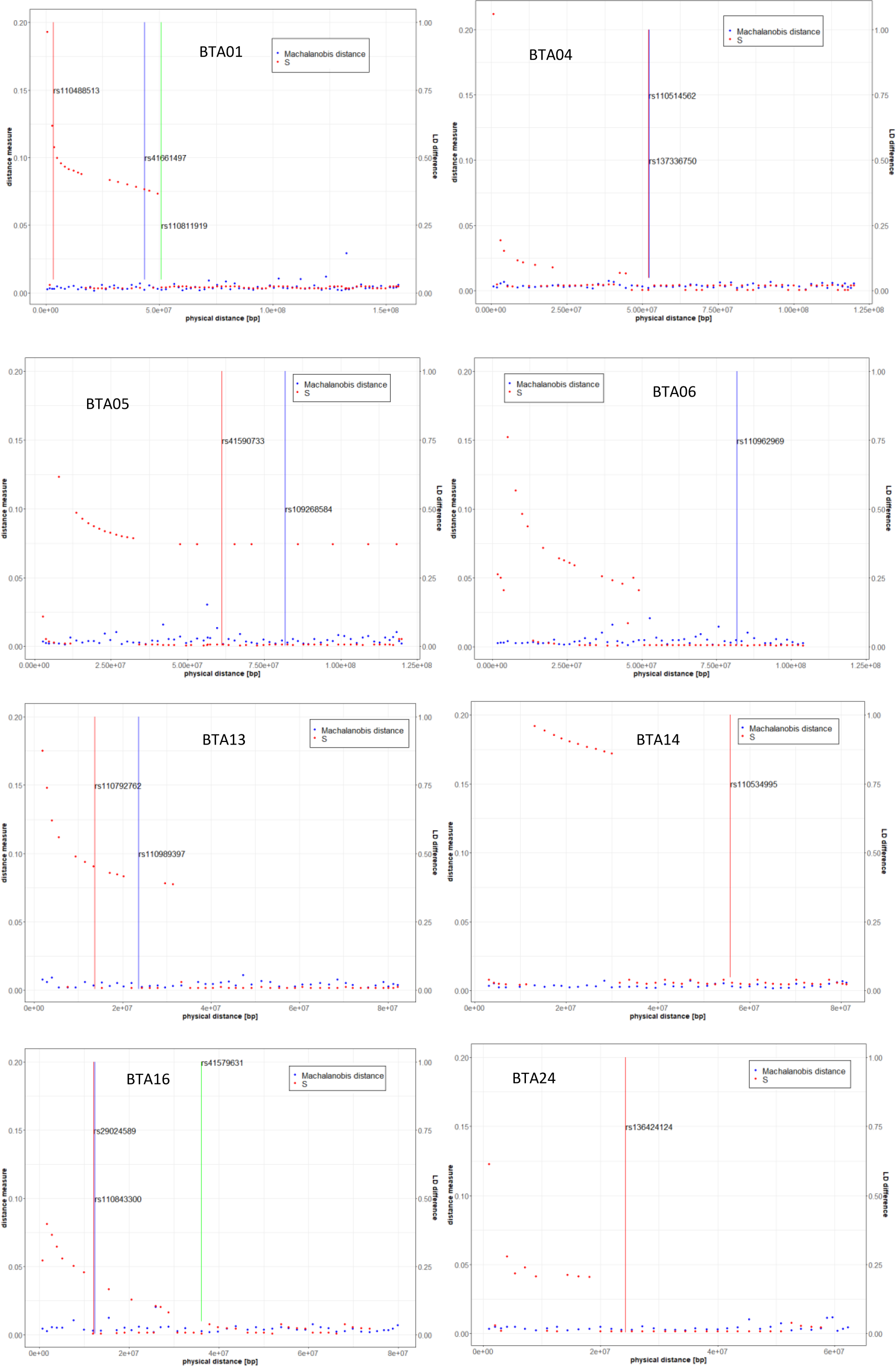
Descriptive statistics of heterogeneity between Braunvieh and Fleckvieh on chromosomes containing SNPs significant for Braunvieh in GWAS. Vertical bars mark the positions of significant SNPs.

Summarising the obtained results, inter-breed differences at some of the 16 significant positions can be explained by inter-breed differences in LD structure (BTA01, BTA05, BTA13 and BTA16) indicated by high values of the S statistics and/or by significant allelic heterogeneity (BTA01, BTA04, BTA05, BTA13 and BTA16). Still polymorphisms on BTA06 (marking SORCS2), BTA14 (marking TMEM74 and EMC2) and BTA24 (marking CCDC178 and KLHL14) are good candidates for Braunvieh-specific associations.

## Discussion

Although the number of published studies related to GWAS for hoof and leg disorders is very limited, their common feature is the lack of overlap in significant results both, between and even within the studies. Similarly, to our study, Wu et al. [9] applied the same GWAS model to feet and leg disorders in three breeds and depending on breed identified different significant regions between Danish Red and Danish Holstein, while no significance was observed in Jersey. In addition, earlier van der Spek et al. [10] found a very low overlap in significance while analysing a cow data set and a bull data set ascertained from the same population of Holstein-Friesian cattle, with only three SNPs in bulls overlapped with 94 SNPs significant for claw disorders in cows. Vargas et al. [11] found no overlap between significant regions defined for binary and categorical feet and leg classification scores in Nelore breed.

Also, in our study we observed no overlap in significance between Braunvieh and Fleckvieh. The potential basis of this phenomenon is either of a technical nature – type I/type II errors due to limited sample size, or of a genetic nature - genetic heterogeneity in the susceptibility to leg diseases between breeds. In humans, Coram et al. [12] reported a similar result regarding loci determining concentration of lipids in blood, where many differences between populations were due to allele frequencies at the candidate SNPs. The same study also points out at the presence of population-specific significant loci, which, as in our study, can be explained by population-specific selection pressure. Another postulated cause of heterogeneity, see e.g. An and Claudianos [2], for their discussion on autism disorder), pointing out at different causal mutations within the common metabolic pathways. Similarly, Wu et al. [9] in the context of hoof and leg disorders in cattle hypothesised that the breed-specific significance hits represent relatively novel mutations, which occurred after breed separation. The third cause of heterogeneity are differences in genetic architecture between breeds, which are manifested by genome-wide (Figure 3), but also by local differences in LD, which were detected in our study within some of the regions harbouring SNPs significant in Braunvieh. Differences in LD patterns were considered in the context of heterogeneity detected between human populations [11].

**Figure 3.**
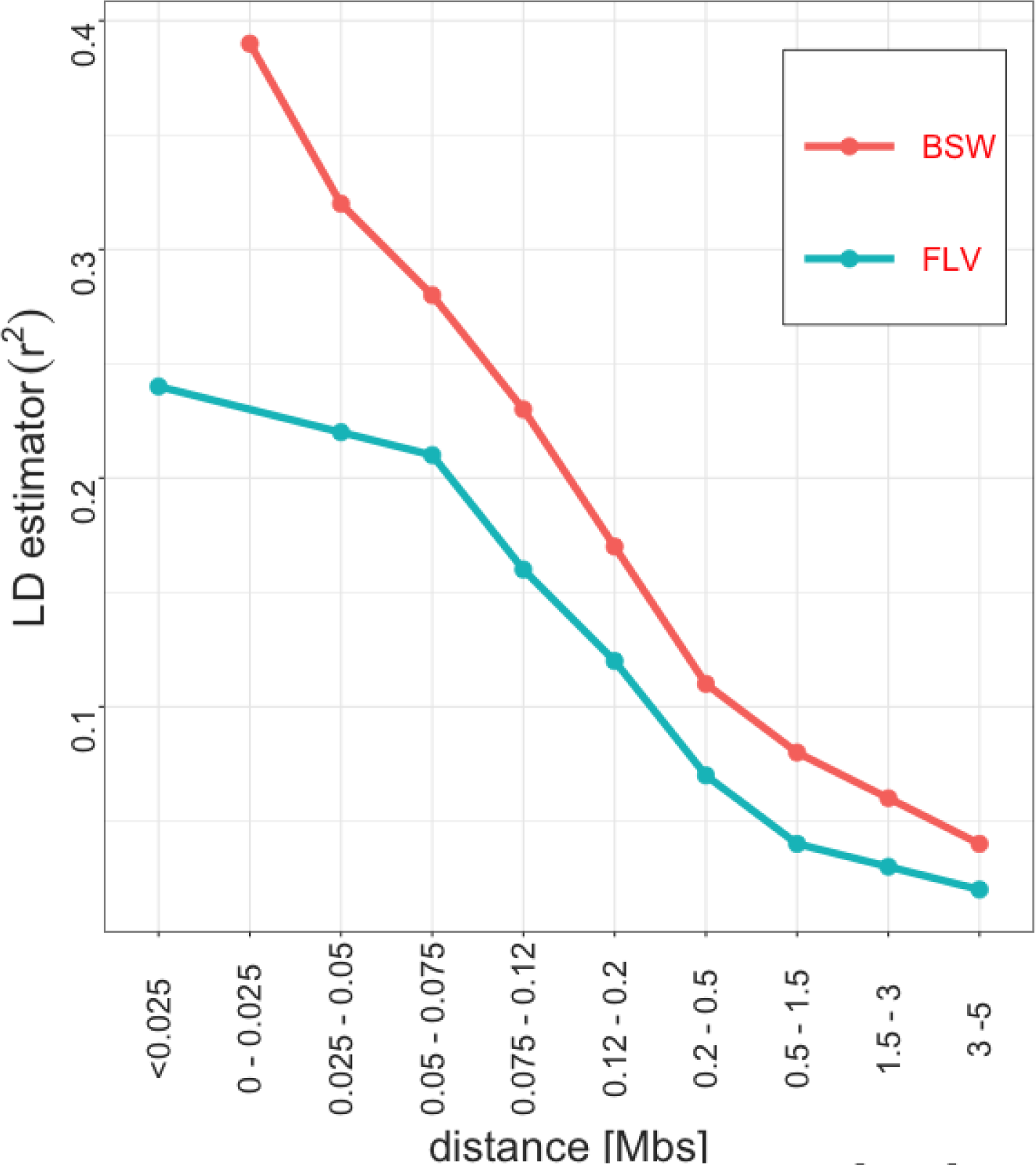
Genome-wide LD decay in Braunvieh and Fleckvieh.

## Conclusions

Based on the current data set it is no possibly to unequivocally confirm/exclude the hypothesis of genetic heterogeneity in the susceptibility to leg disorders between Fleckvieh and Braunvieh. The rationales against the hypothesis comprise: (i) limited power of detection of true associations if the effect size is not large and therefore high rate of spurious associations among detected SNP, (ii) differences in the length of LD blocks, which imply differences in power of detecting the associations, (iii) differences in linkage phase between breeds, which may hamper the detection of causal sites in Fleckvieh or Braunvieh based on the available SNP panel. On the other hand (i) multigenerational differential selection of the breeds – towards dairy production in the case of Braunvieh and towards beef production in the case of Fleckvieh, (ii) no significant allelic heterogeneity, and (iii) no systematic differences in LD structure between the breeds stay in favour of the heterogeneity hypothesis at the significant sites on BTA06, BTA14, and BTA24.

Unfortunately, the data set available for the analysis comprises only common SNPs selected for a commercial microarray, so that we can explore only associations and not the causal mutations, therefore a final verification of the above hypothesis would require a denser SNP map from whole genome sequence.

## Methods

### Dataset

The analyzed data set was collected within the frame of the Efficient Cow project and comprised scores of hoof and leg disorders in Austrian 985 Braunvieh and 1,999 Fleckvieh cows. In particular, the analyzed phenotype comprised the total number of hoof disorders scored until 100^th^ day of lactation. In both breeds, the number of disorders varied between none and five, but the distributions differ with the fraction of diseased cows being higher in Fleckvieh (Figure 1). The cows were genotyped with the GeneSeek^®^ Genomic ProfilerTM HD panel consisting of 76,934 SNPs out of which 74,762 SNPs remained for further analysis after preprocessing based on a minor allele frequency (<0.01) and a per-individual call rate (<99%).

### Genome-wide association study

The genome-wide association study was performed separately for each breed by applying a series of single-SNP mixed linear models implemented in the GCTA software [13] for pseudophenotypes, expressed by cows’ breeding values estimated by Suchocki et al. [14]. For a single SNP the model is given by:

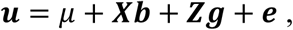

where, ***u*** is a vector of breeding values, *μ* is a general mean, ***b*** is the fixed additive effect of a single SNP, ***X*** is a corresponding design matrix coded as 0, 1 or 2 for a homozygous, heterozygous and the other homozygous genotype respectively, 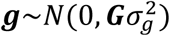 is a random additive polygenic effect with the genomic covariance matrix between cows (***G***) calculated based on SNP genotypes, ***Z*** is an incidence matrix for 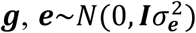 is a residual. The null hypothesis of ***b*** = 0 was tested using the Likelihood Ratio Test with the asymptotic large sample 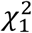 distribution (as implemented into the GCTA). The resulting nominal P-values were transformed into False Discovery Rates [15] to account for multiple testing.

### Analysis of allelic heterogeneity

For each, non-overlapping windows of 50 neighboring SNPs a genomic relationship matrices between cows were calculated, which were then decomposed into principal components, using the PCA subroutine implemented into GCTA [16]. Further on, for each of the windows, differences between the breeds in a 10-dimentional space defined by the first ten eigenvectors (***ε***_**1**_, ***ε***_**2**_, …, ***ε***_**10**_) were quantified using the Machalanobis distance: 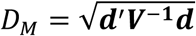. With 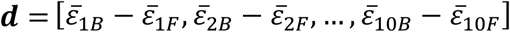 containing differences between averaged eigenvectors for Braunvieh (subscript B) and Fleckvieh (subscript F) and ***V*** representing a pooled covariance matrix of ***ε***_**1**_ and ***ε***_**2**_. The Hotelling test: 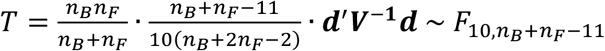 was used to test the null hypothesis of no differences in positions between Braunvieh and Fleckvieh, where *n*_*x*_ is the number of cows representing each breed [17].

The allelic heterogeneity between breeds was tested by calculating the ratio of minor allele frequencies in Fleckvieh (*MAF*_*F*_) and Braunvieh (*MAF*_*B*_) at significant SNP positions. For hypotheses testing, the large sample standard normal distribution of the natural logarithm of the ratio was used: 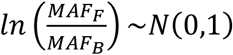.

### Analysis of local linkage disequilibrium patterns

Differences in linkage disequilibrium (LD) patterns between breeds were assessed based on the comparison of LD matrixes constructed for non-overlapping windows of 50 neighboring SNPs. LD between pairs of linked SNPs was quantified using Beagle 4.1 [18], separately for each breed, by the r^2^ coefficient given by 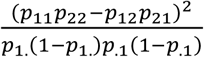, with *p*_*ij*_ corresponding to the frequency of a two-SNP haplotype *ij* ∈ {11,12,21,22}, *p*_1._ and *p*_.1_ representing the marginal SNP allele frequency. Eigenvectors corresponding to the LD matrices were computed separately for each breed, using Python scripts. Inter-breed differences in local linkage disequilibrium were then quantified by:

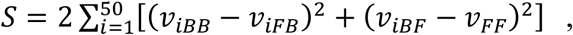

where *ν*_*ijk*_ corresponds the i-th element the 1^st^ principal component vector calculated as the product of the LD matrix of the j-th breed (subscript B for BSW or subscript F for FLV) and the 1^st^ eigenvector of the k-th breed. Following Garcia [19], *S* quantifies differences in the variability of LD between two populations. Furthermore, the genome-wide pattern of linkage disequilibrium (LD) decay with physical distance of pair-wise SNPs were binned into seven types of intervals (0 to 25 kb, 25 to 50 kb, 50 to 100 kb).

## List of abbreviations

BSW: Braunvieh
FDR: false discovery rate
FLV: Fleckvieh
GWAS: genome-wide association study
LD: linkage disequilibrium
PCA: principal component analysis
SNP: single nucleotide polymorphism

## Availability of data and materials

The data that support the findings of this study are available from the ZuchtData EDV-Dienstleistungen GmbH but restrictions apply to the availability of these data, which were used under license for the current study, and so are not publicly available. Data are however available from the authors upon reasonable request and with permission of the ZuchtData EDV-Dienstleistungen GmbH.

## Authors’ contributions

BKS edited data, performed GWAS and heterogeneity analysis. TS edited data and performed some statistical analyses. CEG contributed to writing of the manuscript. HS edited raw data and contributed to writing of the manuscript. MF calculated pairwise linkage disequilibrium. JS provided the concept of the study, performed some statistical analyses and significantly contributed to writing of the manuscript. All authors read and approved the final manuscript.

## Acknowledgements

Computations were carried out at the Poznan Supercomputing and Networking Centre.

